# Night-break photoperiod timing shapes growth trajectories, functional phenotype, and biomass partitioning in *Cannabis sativa* L. in a genotype-dependent manner

**DOI:** 10.64898/2026.09.29.755452

**Authors:** Maria Paula Rodríguez-Roncancio, Laura Camila Castillo-Hernández, Nitro Love, Luz Marina Melgarejo, Felipe Sarmiento

**Affiliations:** Fisiología del estrés y biodiversidad en plantas y microorganismos, Departamento de Biología, Facultad de Ciencias, Universidad Nacional de Colombia, Bogotá, Colombia; Grupo Ingeniería genética de plantas (IGP), Departamento de Biología, Facultad de Ciencias, Universidad Nacional de Colombia, Bogotá, Colombia

**Author notes:** **Corresponding author:** Felipe Sarmiento Salazar, Departamento de Biología, Facultad de Ciencias, Universidad Nacional de Colombia, Bogotá, Colombia, Carrera 30 No. 45-03, Bogotá 111321, Colombia.

**Keywords:** Flowering, physiology, biomass allocation, harvest index, cannabinoids

## Abstract

*Cannabis sativa* L. cultivation in tropical regions or under indoor conditions requires supplemental lighting to maintain plant yields. Night-break (NB) photoperiods offer a cost-effective alternative to conventional day extension, although genotypic responses have not been clearly described. This study evaluated the effects of continuous and pulsed NB treatments on the growth, phenology, physiology, productivity, and cannabinoid accumulation of three *Cannabis sativa* genotypes representing two chemotypes. Plants were exposed to a commercial 18-h photoperiod (T1), a continuous 1-h NB (T2), or four 15-min NB pulses (T3) with equal total duration. Continuous NB produced responses comparable to the commercial photoperiod. In contrast, pulsed NB accelerated phenological progression by two to three weeks and reduced final plant height by 15-28%, depending on genotype, without impairing photosynthetic performance or decreasing floral dry biomass. Harvest index increased under both NB treatments, indicating enhanced biomass allocation to reproductive structures, while cannabinoid accumulation remained primarily genotype-dependent with only minor treatment-specific modulation. These findings demonstrate that NB efficacy depends not only on the duration of nocturnal light but also on its temporal distribution, highlighting temporal organization of nocturnal light as a key regulator of photoperiodic responses and a promising strategy for optimizing cannabis cultivation.

## 1. Introduction

*Cannabis sativa* L. —hereafter referred to as cannabis—is an annual dioecious species cultivated worldwide for medicinal, industrial and pharmaceutical applications (Long et al. 2017; Fordjour et al. 2023) owing to the high-value secondary metabolites synthesized and accumulated in the glandular trichomes of unfertilized female inflorescences (Appendino et al. 2011; Vásquez-Ocmín et al. 2021). Among these metabolites, phytocannabinoids —including Δ -tetrahydrocannabinol (THC), cannabidiol (CBD) and cannabigerol (CBG)—are of particular interest because of their therapeutic and commercial value (Andre et al. 2016; Boggs et al. 2016; Gülck and Møller 2020). Based on cannabinoid composition, cannabis is commonly classified into distinct chemotypes, including CBD-dominant (type III) and CBG-dominant (type IV) plants, which are widely cultivated for medicinal purposes (Salamone et al. 2022). Although cannabinoid accumulation is primarily genetically determined, it is strongly influenced by environmental conditions and floral developmental stage, making the regulation of reproductive development and harvest timing critical for maximizing chemical yield while complying with regulatory thresholds (Aizpurua-Olaizola et al. 2016; Yang et al. 2020; Stack et al. 2021; Chiluwal et al. 2023; Peterswald et al. 2023).

Photoperiod is one of the primary environmental cues regulating seasonal developmental transitions in plants, synchronizing flowering with favorable environmental conditions through the integration of light perception and the circadian clock (Dang et al., 2022; Ishikawa et al., 2005; Pearce et al., 2017; Song et al., 2015). In cannabis, traditionally regarded as a short-day species, short photoperiods promote reproductive development, whereas extended photoperiods maintain vegetative growth and biomass accumulation before the formation of compact floral inflorescences (Zhang et al. 2021; De Prato et al. 2022). However, this classification has been refined by evidence showing that early reproductive events, such as solitary flower formation, occur even under long-day conditions, while short photoperiods induce progressive changes in shoot architecture, including internode shortening and the development of increasingly compact floral structures (van der Werf et al. 1994; Amaducci et al. 2008; Spitzer-Rimon et al. 2019; Trancoso et al. 2022). Photoperiodic responses also vary markedly among genotypes, reflecting differences in flowering thresholds and sensitivity to day length, with highly responsive genotypes exhibiting significant developmental changes following photoperiod adjustments of only a few minutes (Zhang et al. 2021).

Because photoperiod regulates both development and carbon assimilation, alterations in the light–dark cycle may also influence plant physiological performance. Chlorophyll fluorescence parameters, including the effective quantum yield of photosystem II (ΦPSII) and electron transport rate (ETR), together with stomatal conductance (gsw), transpiration and chlorophyll content, provide complementary indicators of photosynthetic function and plant responses to environmental conditions (Farquhar and Sharkey 1982; Genty et al. 1989; Baker and Rosenqvist 2004; Guo and Gan 2005; Paradiso and Proietti 2022). In cannabis, the coordination of photosynthesis, stomatal regulation and light signaling is closely linked to circadian regulation; therefore, photoperiod manipulation may influence developmental processes and changes in photosynthetic performance. (Dang et al. 2022; Venkat and Muneer 2022; Ahsan et al. 2024).

Because photoperiod simultaneously regulates growth, development and physiological performance, these responses should be evaluated as an integrated functional phenotype rather than as isolated traits. Functional phenotypes emerge from the interaction between genotype and the light environment, reflecting how each genotype translates environmental signals into changes in plant architecture, developmental timing, photosynthetic performance and biomass allocation (Violle et al. 2007). Accordingly, traits such as plant height, stem diameter and internode length are better interpreted as developmental trajectories than as final measurements, allowing biologically meaningful comparisons among genotypes and photoperiodic treatments (Archontoulis and Miguez 2015). Likewise, phenological scales such as BBCH describe developmental progression, whereas bud dry weight (BDW) and harvest index (HI) distinguish absolute floral production from the efficiency of biomass partitioning toward reproductive structures (Meier, 1998; Naim-Feil et al., 2021).

Maintaining vegetative growth of cannabis under tropical and subtropical conditions, where day length remains close to 12 h year-long, generally requires supplemental lighting for at least 6 hours to prevent premature flowering, increasing production costs and energy consumption (Spitzer-Rimon et al. 2019; Whipker et al. 2020; Dang et al. 2022). As an alternative, night-break (NB) photoperiods consists of applying light during the dark period to interrupt the perception of a continuous night, thereby modulating photoperiodic development, including floral induction (Higuchi et al. 2012; Cao et al. 2016; Park and Jeong 2020). Although NB treatments have been widely adopted to regulate flowering in horticultural crops, most studies have focused on light duration rather than on the temporal distribution of the nocturnal light signal.

In cannabis, previous studies have shown that NB photoperiods can delay or prevent the development of floral inflorescences while maintaining biomass accumulation at levels comparable to conventional day-extension strategies, although morphological changes, shifts in phenological trajectories, and variation in cannabinoid accumulation have also been reported (Whipker et al. 2020; Gonzalez-Vanegas et al. 2024; Grover et al. 2025). However, existing studies have primarily evaluated NB responses in CBD-dominant genotypes and have focused on the presence or duration of nocturnal light rather than on its temporal structure. Moreover, substantial genotypic variation in photoperiod sensitivity has been documented in cannabis, suggesting that a single lighting strategy is unlikely to be equally effective across contrasting genetic backgrounds (Zhang et al. 2021; Peterswald et al. 2023).

To address this knowledge gap, this study evaluated the effects of two night-break (NB) lighting regimes on the developmental and productive responses of cannabis representing two contrasting chemotypes and three genotypes. Phenological progression, growth trajectories, physiological performance, biomass partitioning, and cannabinoid accumulation were assessed under a commercial day-extension photoperiod, a continuous 1-h NB treatment, and a pulsed NB treatment with the same total duration of nocturnal light. We hypothesized that the effectiveness of NB photoperiods depend not only on the total duration of nocturnal illumination but also on its temporal distribution, and that these responses are modulated by genotype.

## 2. Materials and Methods

### 2.1 Plant material and experimental conditions

Three cannabis genotypes representing two contrasting chemotypes were evaluated. Stellita CBD and Mila are CBD-dominant (type III), whereas Skysun is CBG-dominant (type IV). Four-week-old rooted cuttings were supplied by Panacea CL S.A.S. The experiment was conducted in a commercial greenhouse located in La Vega, Cundinamarca, Colombia (4.74° N, 74.38° W), between December 2024 and March 2025. Plants were spaced 1 m apart and grown under standard Panacea CL S.A management, with a mean air temperature of 30 °C, 61% relative humidity and daily fertigation (1 L daily per plant). Crop nutrition, irrigation and phytosanitary management were maintained under optimal conditions throughout the experiment.

The experiment followed a completely randomized factorial design with genotype and photoperiod treatment as fixed factors. Individual plants were considered the experimental units, and repeated measurements were performed throughout the experimental period. Each genotype × treatment combination consisted of 26 plants distributed across two independent cultivation beds separated by black polyethylene curtains (7-mm thickness) to prevent light contamination among treatments, whereas genotypes were randomly distributed within each treatment compartment.

### 2.2 Photoperiod treatments and light characterization

The experimental cycle lasted 14 weeks, comprising six weeks of vegetative growth under the assigned photoperiod treatments followed by eight weeks under an inductive 12 h light/12 h dark photoperiod until harvest (Fig. 1). LED lamps were installed 2 m above the plant canopy. Three photoperiod treatments were evaluated: T1, a commercial day-extension photoperiod (control) consisting of 18 h of light (natural daylight from 06:00 to 18:00, followed by artificial lighting from 18:00 to 00:00) and 6 h of darkness; T2, a continuous 1-h night-break (NB) treatment comprising 12 h of daylight (06:00–18:00), 5 h of darkness and 1 h of artificial light (23:00–00:00); and T3, a pulsed NB treatment consisting of the same total duration of nocturnal light (1 h), delivered as four 15-min light pulses separated by 45 min of darkness (20:45–21:00, 21:45–22:00, 22:45–23:00 and 23:45–00:00).

**Fig 1.**
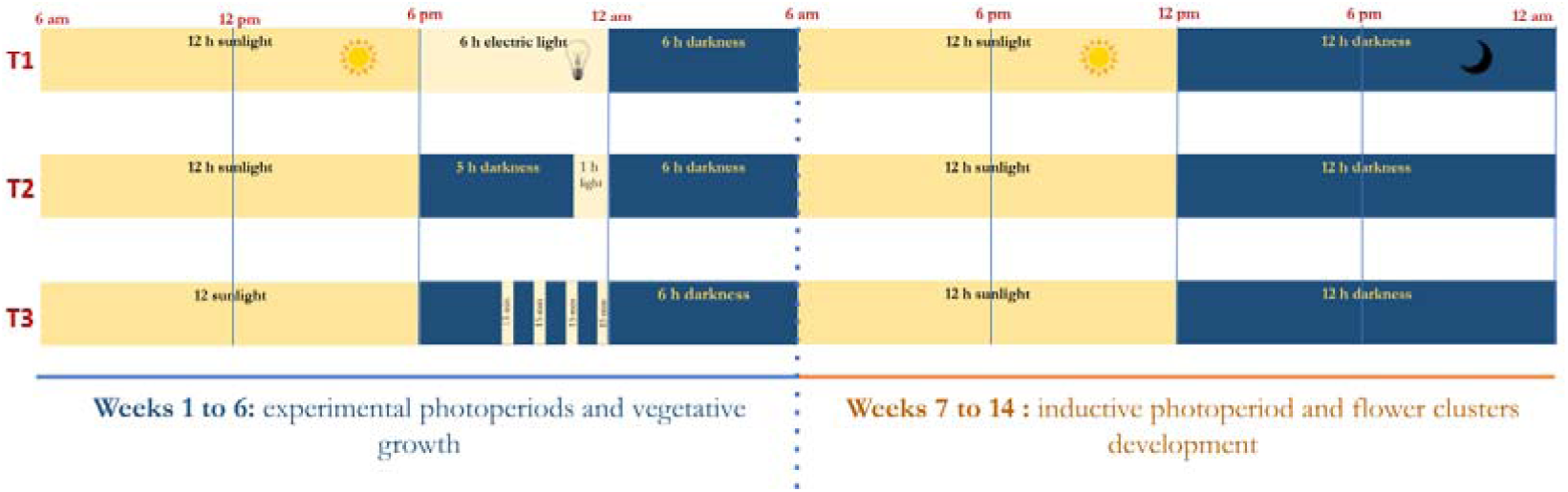
Experimental design and photoperiod schedules applied to cannabis during the 14-week cultivation cycle. Plants were maintained under three photoperiod treatments during the six-week vegetative phase, followed by an inductive 12 h light/12 h dark photoperiod for eight weeks until harvest. The diagram illustrates the light and dark schedules for each treatment and the transition to the flowering-inductive photoperiod at week 7.

Incident radiation was characterized at canopy level (approximately 45 cm above the substrate) using a LI-180 spectroradiometer (LI-COR Biosciences, Lincoln, NE, USA). Daytime photosynthetic photon flux density (PPFD) ranged from 300 to 800 μmol m ² s ¹ with a balanced spectral distribution (R:FR = 1.1-1.4). Night-break lighting was provided by 20-W, 6500 K LED lamps, delivering a PPFD between 3.9 and 8.8 μmol m ² s ¹ with spectra enriched in blue and red wavelengths and minimal far-red contribution (R:FR ≈ 10.9).

### 2.3 Morphological, phenological and physiological measurements

#### Growth measurements

Six plants were randomly selected from each genotype × treatment combination for repeated measurements at weeks 3, 5, 6, 9, 11 and 14. Plant height, stem diameter and apical internode length were measured throughout the experiment. Plant height was recorded from the stem base to the apical meristem using a measuring tape, whereas stem diameter and apical internode length were measured with a digital Vernier caliper.

#### Phenological assessment

Phenological development was assessed using the BBCH scale (Hess et al. 1997; Meier 1998), adapted for cannabis by Gonzalez-Vanegas et al. (2024). Stages 5 (inflorescence emergence) to 6 (flowering with visible pistils) were recorded. A developmental stage was assigned when at least 50% + 1 of the plants within each genotype × treatment combination reached that stage.

#### Physiological measurements

Total stomatal conductance (gsw), effective quantum yield of photosystem II (ΦPSII), electron transport rate (ETR) and transpiration (E) were measured using an LI-600 fluorometer–porometer (LI-COR Biosciences, Lincoln, NE, USA) between 09:00 and 10:00 h on the central leaflet of a fully expanded leaf located in the middle third of each plant (Gonzalez-Vanegas et al. 2024). Relative chlorophyll content was estimated using a SPAD-502 meter (Konica Minolta, Japan) from five readings per selected leaf. SPAD values were converted to total chlorophyll content using a calibration curve generated from pigment extraction in 80% acetone and spectrophotometric absorbance at 664, 649 and 470 nm (Lichtenthaler 1987; Melgarejo et al. 2010; Lozano-Montaña et al. 2021).

### 2.4 Harvest measurements and cannabinoid analysis

At the end of the experiment (week 14), destructive harvest was performed on representative branches collected from three plants per genotype × treatment combination. The sampling strategy was based on biomass allometric partitioning and canopy structural representativeness, considering the vertical architectural heterogeneity of cannabis (Poorter et al. 2012; Danziger and Bernstein 2021). Plants were separated into leaves, stems, branches and inflorescences, and fresh biomass of each organ was recorded. Leaves, stems and branches were oven-dried at 65 °C to constant weight (5 days), whereas inflorescences were dried at 65 °C for 24 h (Gonzalez-Vanegas et al. 2024). Prior to weighing, non-glandular material was removed from the inflorescences. Bud dry weight (BDW) was determined as the dry mass of marketable floral tissue. Harvest index (HI) was calculated as the ratio of inflorescence dry weight to total plant dry weight, and values were extrapolated according to the number of branches per plant.

For cannabinoid analysis, 2-g subsamples of dried inflorescences from the upper canopy were collected from three plants per genotype × treatment combination. Cannabinoid concentrations were determined by high-performance liquid chromatography with photodiode array detection (HPLC-PDA) following the accredited PAL OT035 V3 method (ISO/IEC 17025:2017; LIAC Laboratory, Universidad de La Salle, Colombia). Total CBD, CBG and THC contents (g 100 g□¹ dry flower) were calculated by stoichiometric conversion of their acidic forms to neutral cannabinoids.

### 2.5 Statistical analysis

All statistical analyses were performed in RStudio (version 2025.05.1). Physiological variables were analyzed independently for each sampling week using analysis of variance (ANOVA) when assumptions of normality and homogeneity of variance were met, or Kruskal–Wallis tests when assumptions were violated. Significant differences among treatments were evaluated using Tukey’s honestly significant difference (HSD) test or Dunn’s test, respectively (α = 0.05). Phenological development (BBCH scale) was analyzed as longitudinal ordinal data using the F2-LD-F1 model implemented in the nparLD package (Noguchi et al. 2012), considering genotype, treatment and time as experimental factors. BBCH scores were rescaled to a normalized Adjusted Phenological Stage (APS) ranging from 0 to 1 to facilitate statistical comparisons. Pairwise post hoc comparisons were performed using Tukey-adjusted contrasts. Time to each BBCH developmental stage was expressed as days after planting (DAP), calculated as the number of days elapsed from transplanting to the first observation of each phenological stage. DAP was summarized for each genotype × treatment combination as the median, interquartile range (IQR), and the proportion of plants reaching each stage.

Growth trajectories were modeled using nonlinear regression. Plant height was described using a three-parameter logistic model (Asym, xmid and scal), stem diameter using a Michaelis–Menten model (Asym and K), and apical internode shortening using a negative exponential model (parameters a and b) (Archontoulis and Miguez 2015). The model equations and their biological interpretation are provided in Supplementary Table 1.

Multivariate responses were explored by principal component analysis (PCA), followed by permutational multivariate analysis of variance (PERMANOVA) based on Euclidean distances with 999 permutations, evaluating the effects of genotype, treatment, sampling week and their interactions.

## 3. Results

### 3.1 Growth trajectories

Plant growth followed nonlinear trajectories that were adequately described by the selected models (Fig. 2; Table 1). Plant height exhibited a sigmoidal growth pattern fitted by the three-parameter logistic model, whereas stem diameter and apical internode length were described by Michaelis-Menten and negative exponential models, respectively. Estimated model parameters are summarized in Table 1.

**Fig 2.**
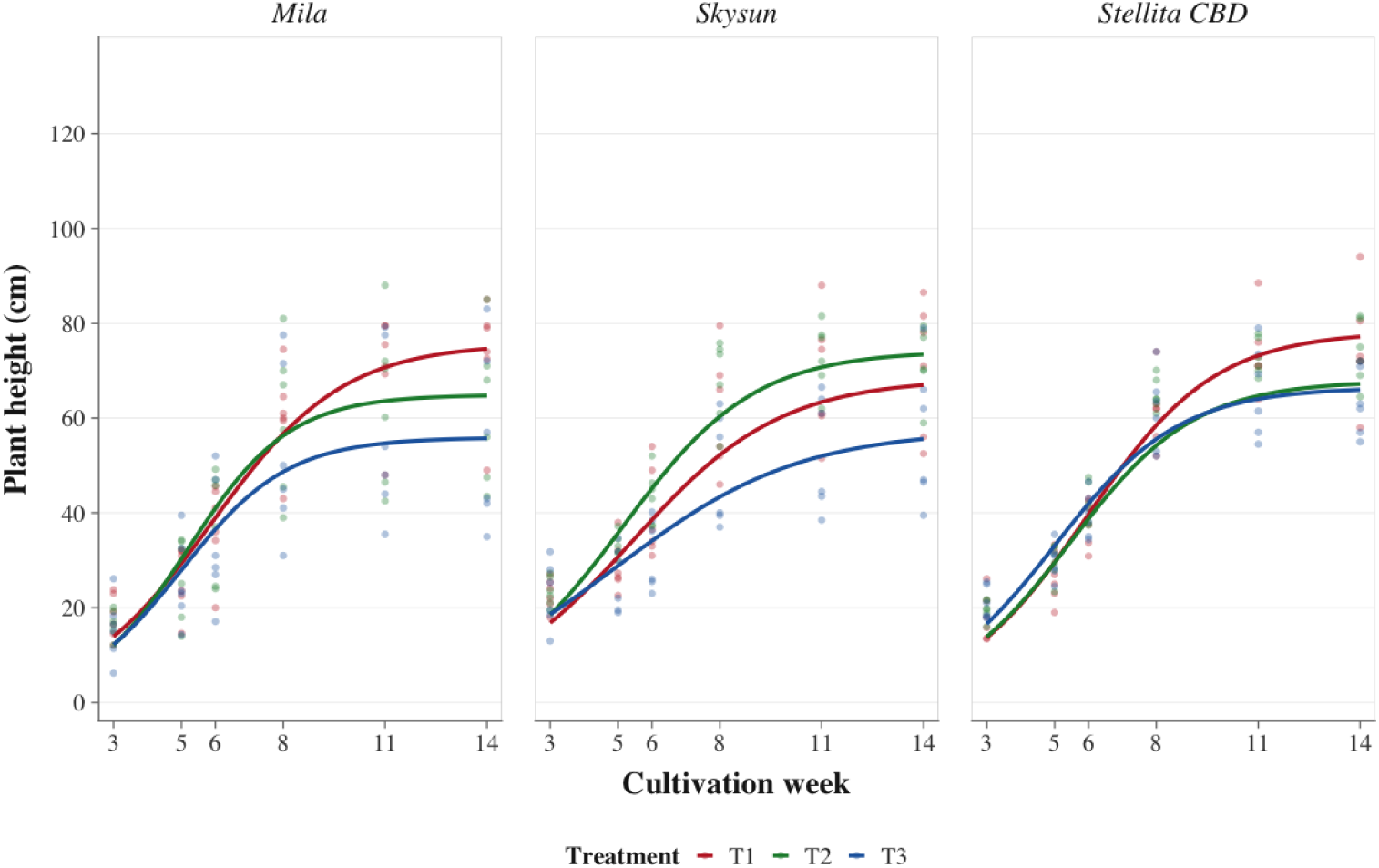
Fitted logistic mixed-effects models describing plant height trajectories of three *Cannabis sativa* genotypes (MI, Mila; SS, Skysun; ST, Stellita CBD) under three photoperiod treatments (T1, commercial day extension; T2, continuous 1-h night break; T3, pulsed night break, 4 × 15 min). Lines represent the fitted model predictions and points correspond to the observed data. Stem diameter was less responsive to photoperiod treatments than plant height. No significant treatment effects were detected for either the asymptotic diameter (Asym) or the growth-rate parameter (K) in Mila or Skysun. In contrast, Stellita CBD exhibited a significantly greater maximum stem diameter under T1 than under T3, while T2 showed intermediate values. A similar pattern was observed for K, indicating a slower progression toward maximum stem diameter under the pulsed NB treatment.

**Table 1.**
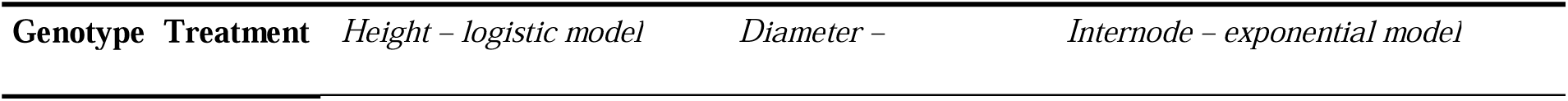
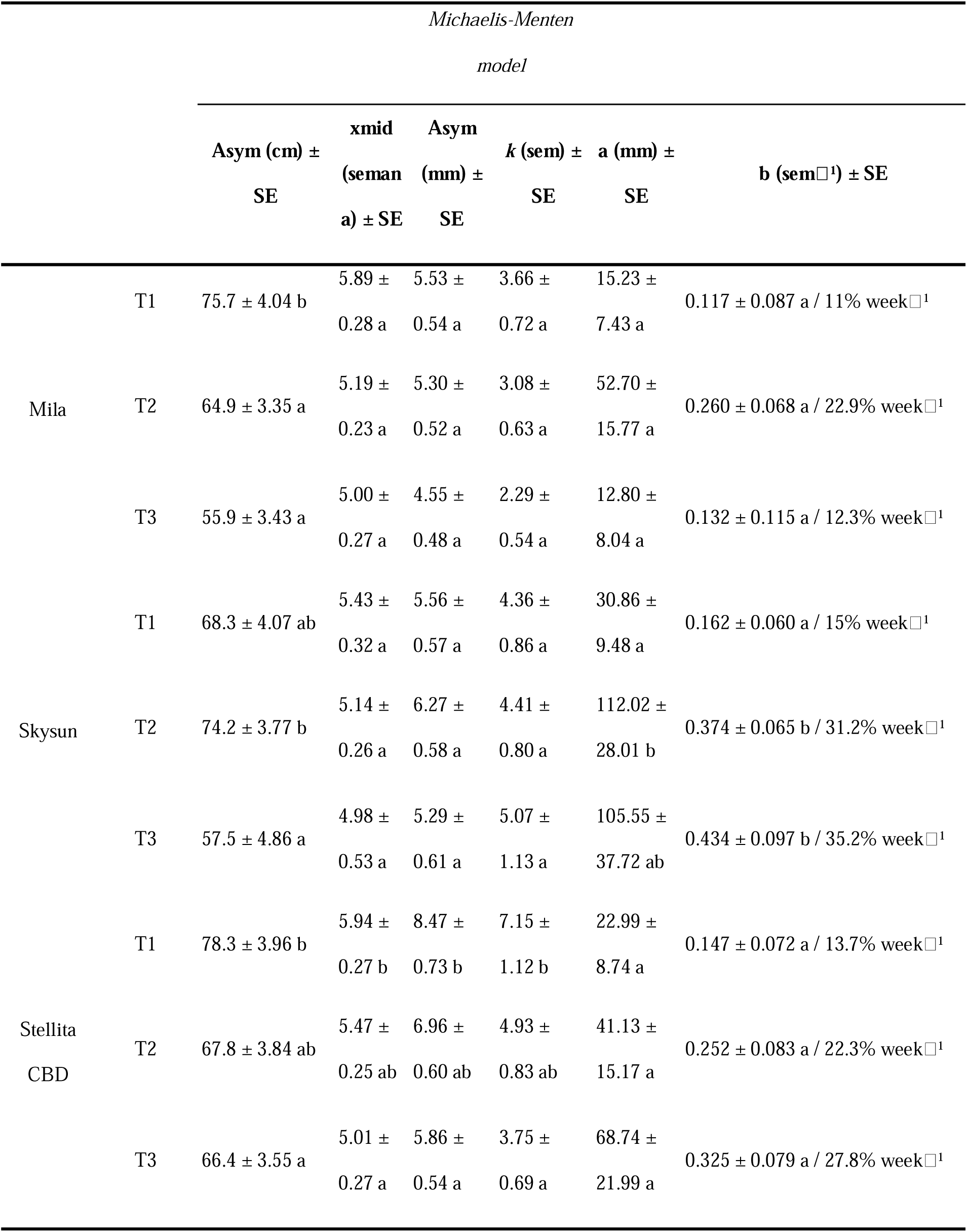
Estimated parameters of nonlinear growth models fitted for plant height, stem diameter and apical internode length in three cannabis genotypes subjected to three photoperiod treatments. Values are presented as mean ± standard error (SE). **Asym**, estimated asymptotic maximum value; **xmid**, time (weeks) to reach 50% of the asymptotic plant height; **K**, diameter growth-rate constant; **a**, estimated initial internode length; **b**, exponential rate of internode shortening. T1, commercial day extension (control); T2, continuous 1-h night break; T3, pulsed night break (4 × 15 min). Different letters within each genotype and parameter indicate significant differences among treatments according to post hoc comparisons (P < 0.05).

Night-break treatments differentially affected the estimated maximum plant height (Asym) depending on genotype. The pulsed NB treatment (T3) consistently reduced asymptotic plant height across all three genotypes, whereas the continuous NB treatment (T2) generally produced values comparable to the commercial day-extension treatment (T1). The reduction was most pronounced in Mila, where T3 decreased the estimated maximum height by approximately 26% relative to the control. In Skysun, the greatest asymptotic height was observed under T2, whereas Stellita CBD exhibited significantly lower values under T3 than under T1 (Table 1). The parameter *xmid*, representing the time required to reach 50% of the estimated maximum height, differed significantly only in Stellita CBD, where plants under T3 reached the inflection point approximately one week earlier than those under the control treatment. No significant differences among treatments were detected in Mila or Skysun. Likewise, the parameter *scal*, describing the slope of the logistic curve around the inflection point, was not significantly affected by photoperiod treatment in any genotype.

Apical internode length decreased progressively throughout the experimental period, consistent with the transition from vegetative to reproductive development. The estimated initial internode length (a) differed significantly only in Skysun, where plants under T2 exhibited greater initial values than those under the control. More importantly, the parameter b, representing the rate of internode shortening, also differed significantly only in Skysun, with both NB treatments producing faster internode shortening than the control. In Mila and Stellita CBD, although parameter b tended to increase under NB treatments, these differences were not statistically significant.

### 3.2 Phenological development

Phenological progression was assessed across the principal developmental stages of cannabis, from vegetative biomass accumulation to the development of pistillate floral inflorescences (Supplementary Table 2) and was significantly affected by genotype, photoperiod treatment and time (P < 0.001). Significant genotype × treatment × time interactions indicated that the effect of photoperiod depended on the genetic background (Fig. 3).

**Fig 3.**
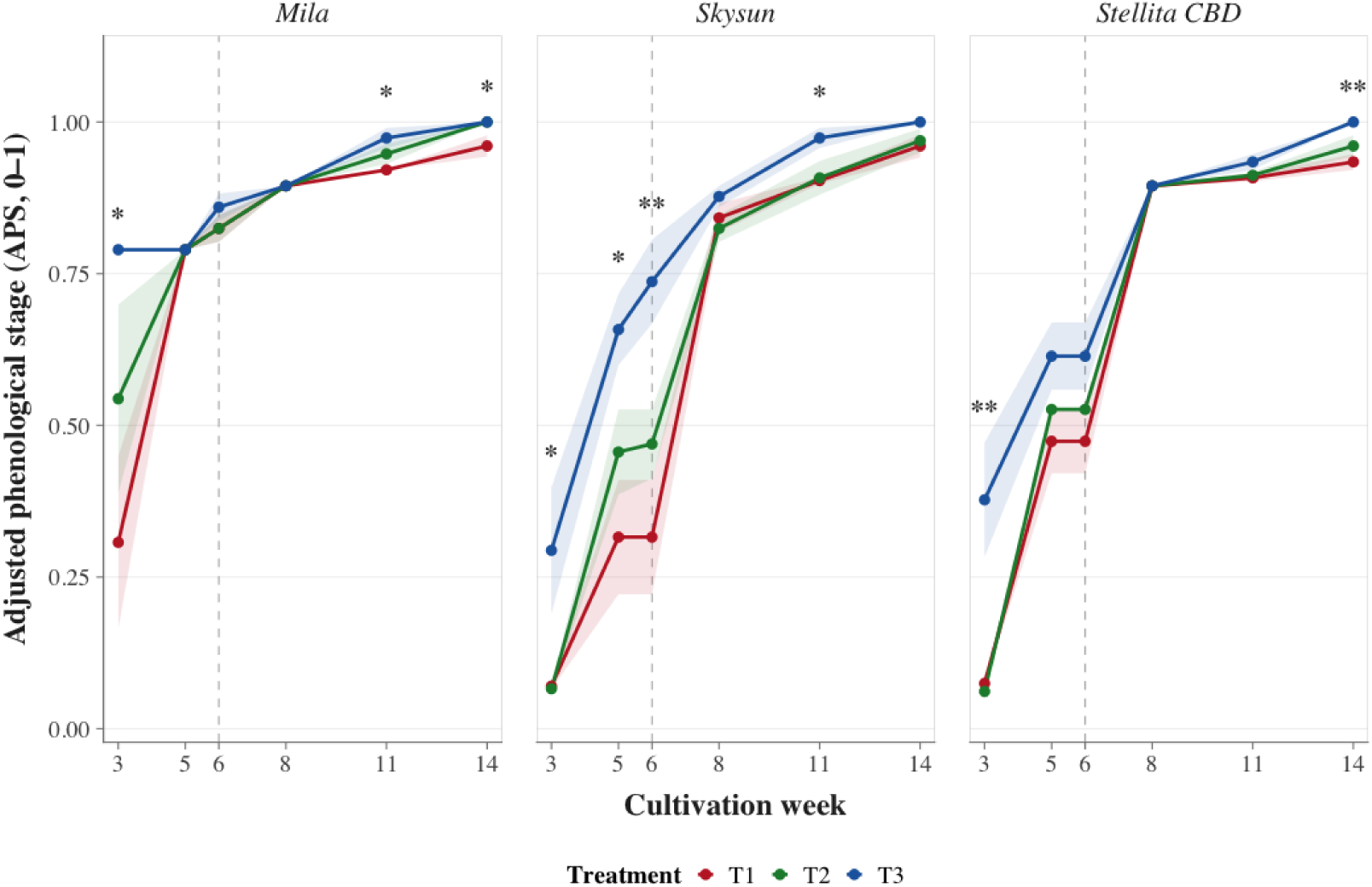
Adjusted phenological progression (Adjusted Phenological Stage, APS) of three Cannabis sativa genotypes (MI, Mila; SS, Skysun; ST, Stellita CBD) under three photoperiod treatments (T1, commercial day extension; T2, continuous 1-h night break; T3, pulsed night break, 4 × 15 min). APS represents a normalized BBCH scale ranging from 0 (vegetative stage) to 1 (most advanced stage of pistillate inflorescence development measured). Asterisks indicate sampling weeks in which significant differences among treatments were detected within each genotype according to nonparametric post hoc comparisons (P < 0.05).

Overall, the pulsed night-break treatment (T3) accelerated phenological development relative to the commercial day-extension treatment (T1), although the magnitude and timing of this response varied among genotypes. In contrast, the continuous night-break treatment (T2) generally followed a phenological trajectory comparable to that of the control throughout the experimental period. Genotype-specific responses were evident in the weekly post hoc comparisons (Fig. 3). In Mila, significant differences were detected only at specific sampling dates, with T3 showing more advanced phenological development than T1 during weeks 3 and 11. By week 14, APS values under T2 and T3 were significantly greater than those under T1, whereas no differences were detected between the two NB treatments. Skysun exhibited the strongest response to photoperiod manipulation, with T3 consistently producing higher APS values than T1 during weeks 3, 5, 6 and 11, and exceeding T2 during week 3. In Stellita CBD, T3 accelerated phenological progression during the early stages of development, showing significantly higher APS values than both T1 and T2 at week 3, whereas differences disappeared during intermediate stages and reappeared at harvest, when T3 again exceeded the control.

Days after planting (DAP) further supported these patterns (Supplementary Table 3). Across the three genotypes, T3 reduced the time required to reach reproductive BBCH stages by approximately 7 to 21 days relative to the control. The greatest acceleration occurred in Skysun, where reproductive stages (BBCH 51, 61 and 69) were reached 17–21 days earlier under T3 than under T1. Stellita CBD showed earlier progression through intermediate stages (BBCH 35-51), whereas Mila exhibited smaller but consistent advances during later reproductive stages (BBCH 61-69). Collectively, these results demonstrate that the temporal distribution of nocturnal light accelerated reproductive development in a genotype-dependent manner, whereas continuous NB produced phenological responses largely comparable to those observed under the commercial day-extension treatment.

### 3.3 Productivity and total cannabinoid concentrations

Bud dry weight (BDW) was significantly affected by genotype, photoperiod treatment, and their interaction (P < 0.001; Fig. 4A). However, genotype explained a larger proportion of the variation than photoperiod treatment. In Skysun, both NB treatments increased BDW relative to the control, although these differences were not statistically significant. Stellita CBD exhibited moderate, non-significant reductions under both NB treatments. In contrast, Mila showed significantly greater BDW under the commercial day-extension treatment (T1) than under either continuous (T2) or pulsed (T3) NB, corresponding to reductions of approximately 46% and 44%, respectively.

**Fig 4.**
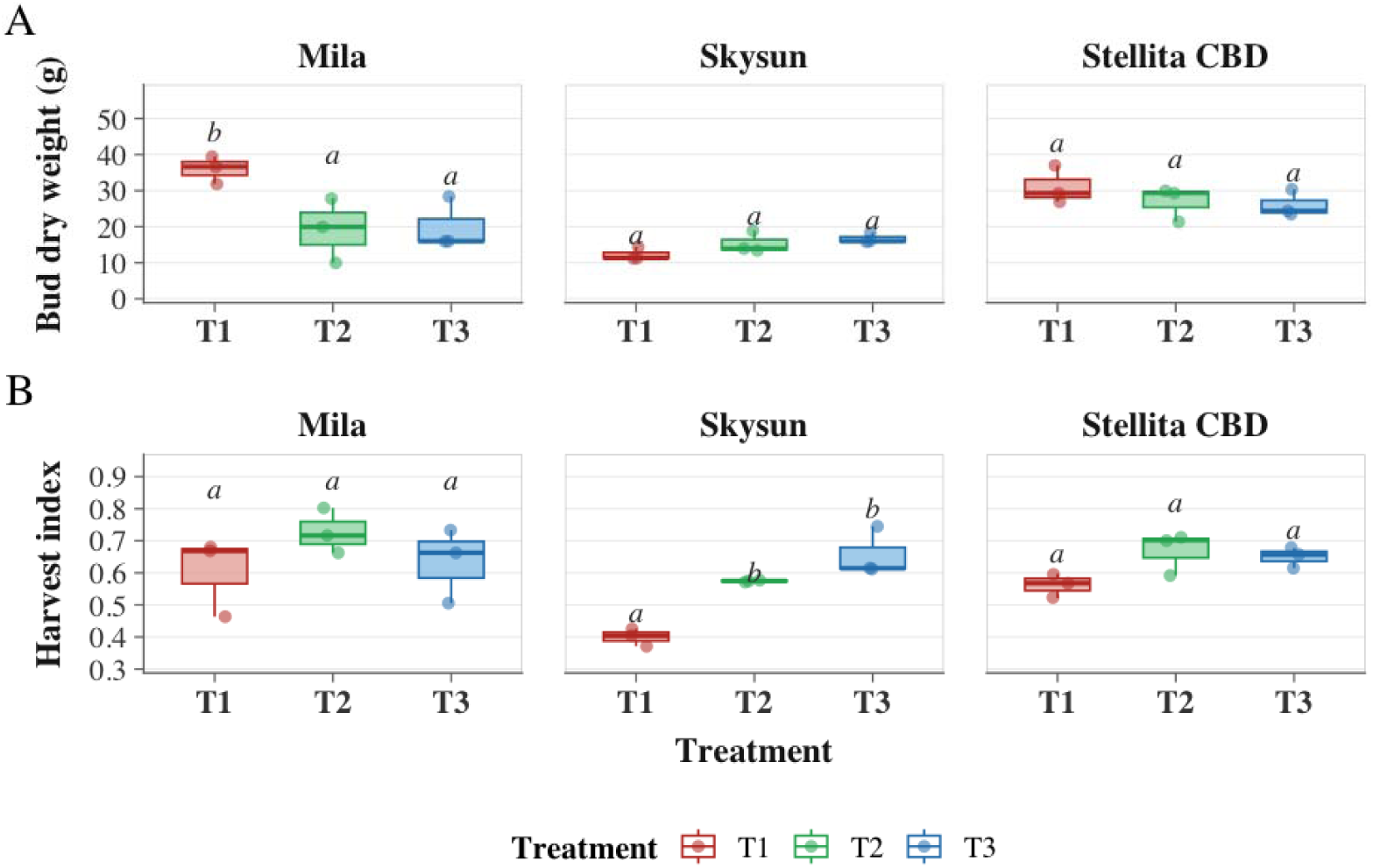
Boxplots showing bud dry weight (BDW; A) and harvest index (HI; B) in three *Cannabis sativa* genotypes subjected to three photoperiod treatments: T1, commercial day extension (control); T2, continuous 1-h night break; and T3, pulsed night break (4 × 15 min). Boxes represent the interquartile range, the central line indicates the median, and whiskers represent data dispersion. Different letters indicate significant differences among treatments within each genotype according to Tukey’s post hoc test (*P* < 0.05).

Harvest index (HI) differed significantly among genotypes (P = 0.017) and photoperiod treatments (P < 0.001), whereas the genotype × treatment interaction was not significant, indicating a consistent treatment effect across genotypes (Fig. 4B). Overall, both NB treatments increased HI relative to the control. The largest response occurred in Skysun, where HI increased by approximately 44% under T2 and 64% under T3 compared with T1. Stellita CBD and Mila exhibited similar trends, although differences among treatments were not statistically significant.

Total cannabinoid concentrations at harvest were primarily determined by genotype (P < 0.001 for CBD, THC, and CBG), whereas photoperiod acted as a secondary, genotype-dependent factor (Table 2). Significant genotype × treatment interactions were detected for CBD and THC, whereas CBG was significantly affected by both genotype and treatment.

**Table 2.**
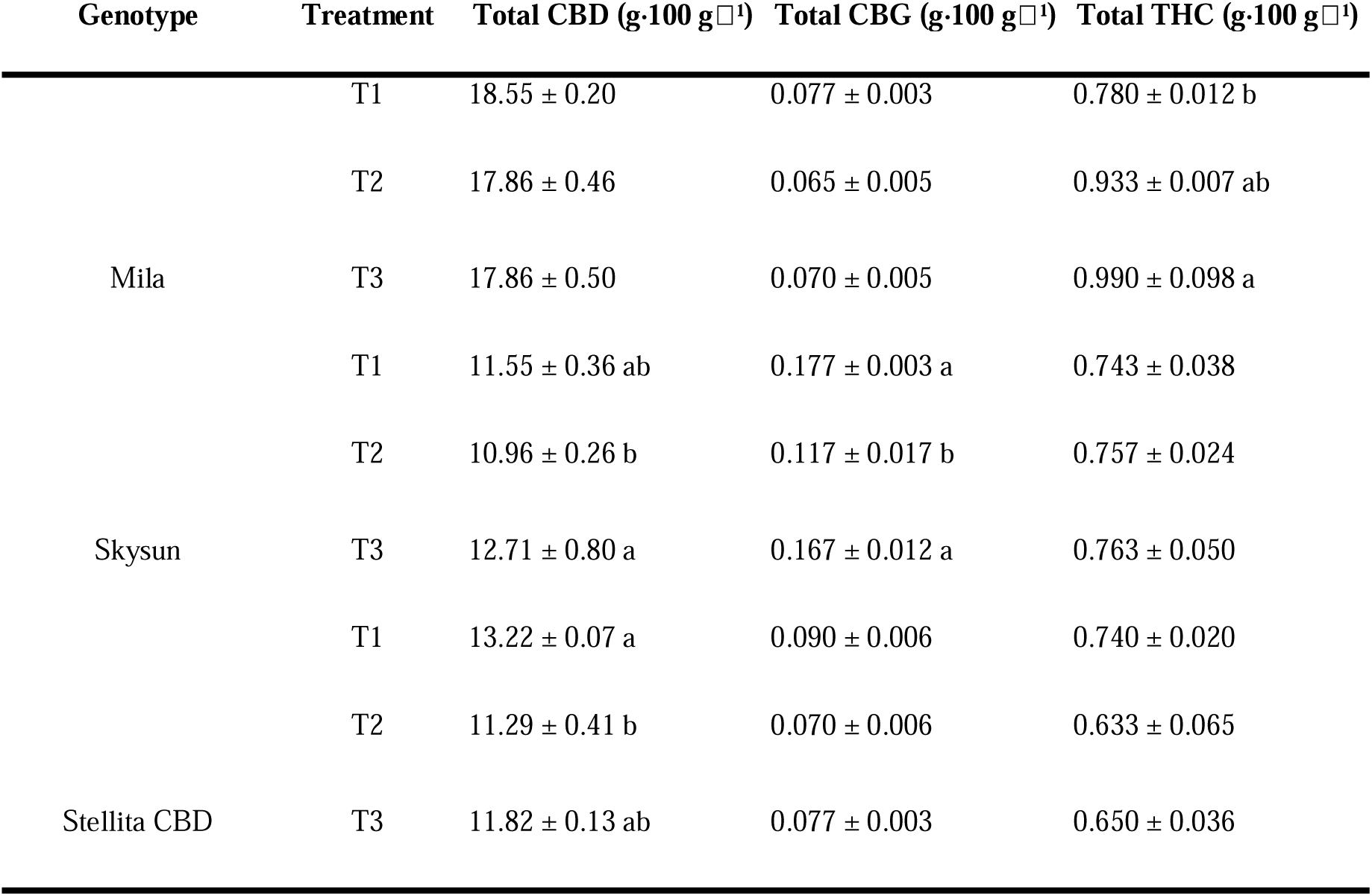
Total cannabinoid concentrations in dried inflorescences of three cannabis genotypes subjected to three photoperiod treatments. Values are presented as mean ± standard error (SE) and expressed as g 100 g ¹ dry biomass. T1, commercial day extension (control); T2, continuous 1-h night break; T3, pulsed night break (4 × 15 min). Different letters within each genotype indicate significant differences among treatments according to Tukey’s post hoc test (*P* < 0.05).

CBD responses varied among genotypes. In Skysun, T3 produced higher CBD concentrations than T2, whereas Stellita CBD showed significantly lower CBD concentrations under T2 than under the control, with T3 displaying intermediate values. In Mila, CBD concentrations remained similar across treatments. THC concentration was largely stable across treatments, although a significant increase under T3 was observed in Mila relative to the control. No significant treatment effects were detected in either Skysun or Stellita CBD. CBG accumulation was highest in Skysun and represented the cannabinoid most responsive to photoperiod treatment. Continuous NB (T2) significantly reduced CBG concentration relative to both the control and the pulsed NB treatment (T3). In contrast, Mila and Stellita CBD maintained low CBG concentrations regardless of photoperiod treatment.

### 3.4 Physiological responses

Physiological performance remained remarkably stable throughout the crop cycle despite differences in photoperiod treatments (Fig. 5). Effective quantum yield of photosystem II (Φ_PSII_), stomatal conductance (gsw), and chlorophyll content exhibited genotype- and time-dependent variation, but no evidence of sustained physiological impairment associated with either night-break treatment was observed. Environmental conditions recorded during the experiment are presented in Supplementary Figure 1. Effective quantum yield of PSII (Φ_PSII_) remained within the expected range for photosynthetically active leaves (0.65–0.75) throughout the experiment (Fig. 5A). Minor temporal fluctuations were observed across genotypes, with lower values during week 3 followed by a gradual increase and a slight decline toward harvest. No consistent treatment effects were detected. Stomatal conductance (gsw) exhibited greater temporal variability than ΦPSII (Fig. 5B). During week 8, Skysun and Stellita CBD displayed significantly greater gsw under NB treatments than under the control, whereas by harvest (week 14) this pattern was reversed, with higher conductance under T1, particularly in Mila and Stellita CBD. These responses were genotype dependent and no consistent treatment effect was maintained throughout the crop cycle.

**Fig 5.**
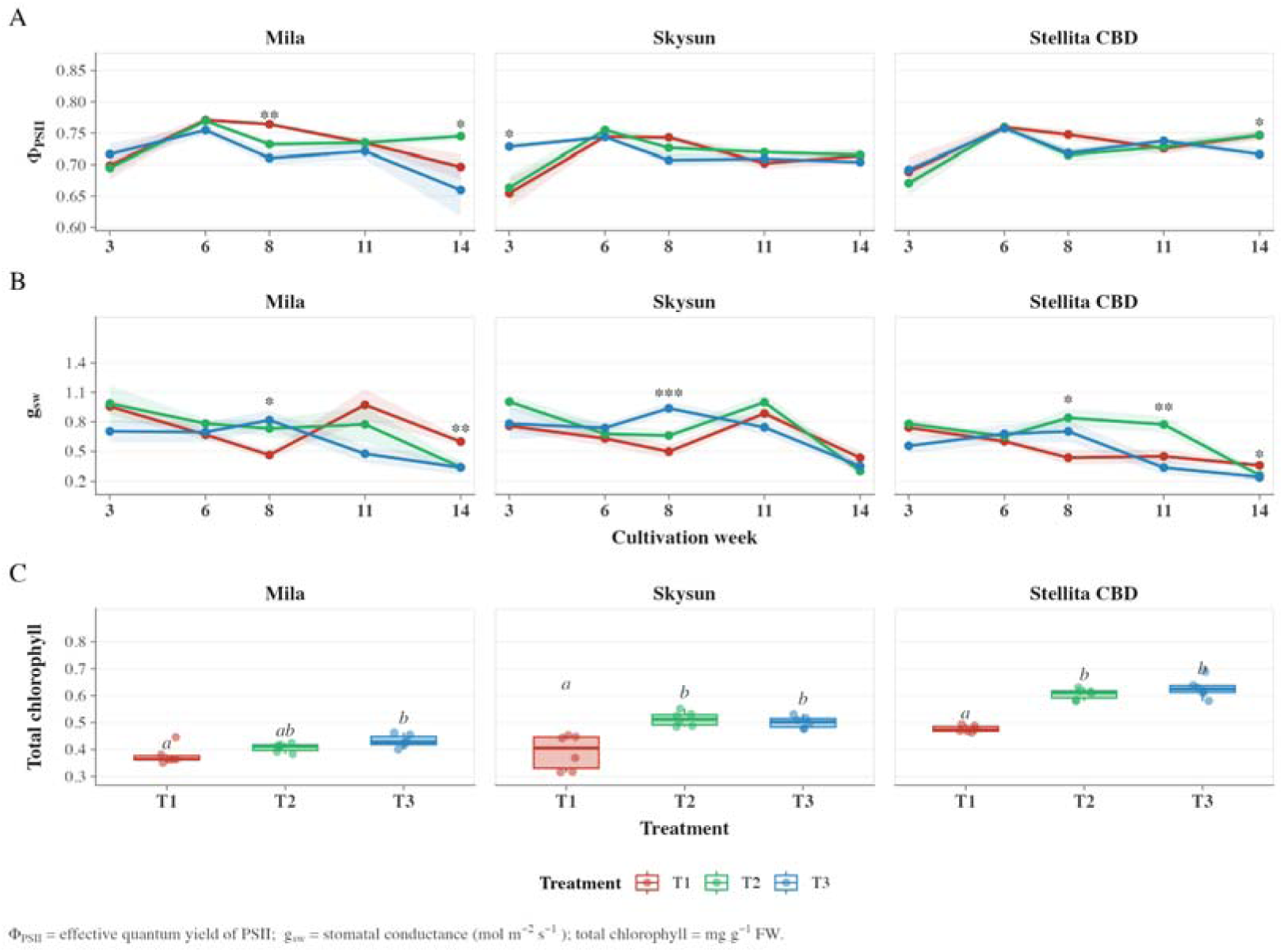
Physiological responses of three *Cannabis sativa* genotypes under three photoperiod treatments (T1, commercial day extension; T2, continuous 1-h night break; T3, pulsed night break, 4 × 15 min). (A) Effective quantum yield of photosystem II (Φ_PSII_). (B) Stomatal conductance (gsw). (C) Total chlorophyll content at week 11, corresponding to the sampling date showing the greatest divergence among treatments. Points represent means and shaded areas indicate the standard error. Asterisks denote significant differences among treatments within each genotype and sampling week (P < 0.05; *P < 0.01).

Total chlorophyll content showed genotype-dependent variation and a marked influence of photoperiod treatment at specific stages of the crop cycle (Supplementary Figure 2). Under T3, and to a lesser extent T2, chlorophyll concentrations were consistently higher than T1 from weeks 8 to 11 across most genotypes (Fig. 5C). In Skysun, significant differences were detected at weeks 8, 11 and 14 (P < 0.05 in all cases), with T3 consistently showing the highest values. In Stellita CBD, treatment effects emerged from week 8 and were most pronounced at week 11, where both T3 and T2 exhibited significantly higher chlorophyll concentrations than T1 (P < 0.05). In Mila, significant differences among treatments were detected only at week 11, with T3 presenting higher chlorophyll concentrations than T1.

Collectively, these results indicate that photoperiod manipulation altered developmental trajectories without causing persistent impairment of photosynthetic performance.

### 3.5 Multivariate analysis

Principal component analyses (PCA) performed throughout the crop cycle revealed progressive changes in the multivariate relationships among morphological and physiological traits (Supplementary Figure S3). The highest proportion of explained variance was observed at week 8 (68.9%), indicating the period of greatest multivariate differentiation among genotype × treatment combinations. The variables contributing to the principal components also shifted throughout plant development. At weeks 6 and 8, PC1 was primarily associated with physiological traits (ETR, gsw, and chlorophyll content) together with plant size (height and stem diameter), whereas PC2 was mainly explained by ΦPSII. By week 11, PC1 was dominated by architectural traits, while PC2 primarily reflected physiological variation. At harvest (week 14), shown in Fig. 6, the first two principal components explained 63.8% of the total multivariate variation. PC1 described a gradient of plant vigor and productivity, characterized by positive contributions from plant height, stem diameter, bud dry weight (BDW), chlorophyll content, and stomatal conductance. In contrast, PC2 was primarily associated with harvest index (HI) and Φ_PSII_, reflecting variation in biomass partitioning and photosynthetic performance.

**Fig 6.**
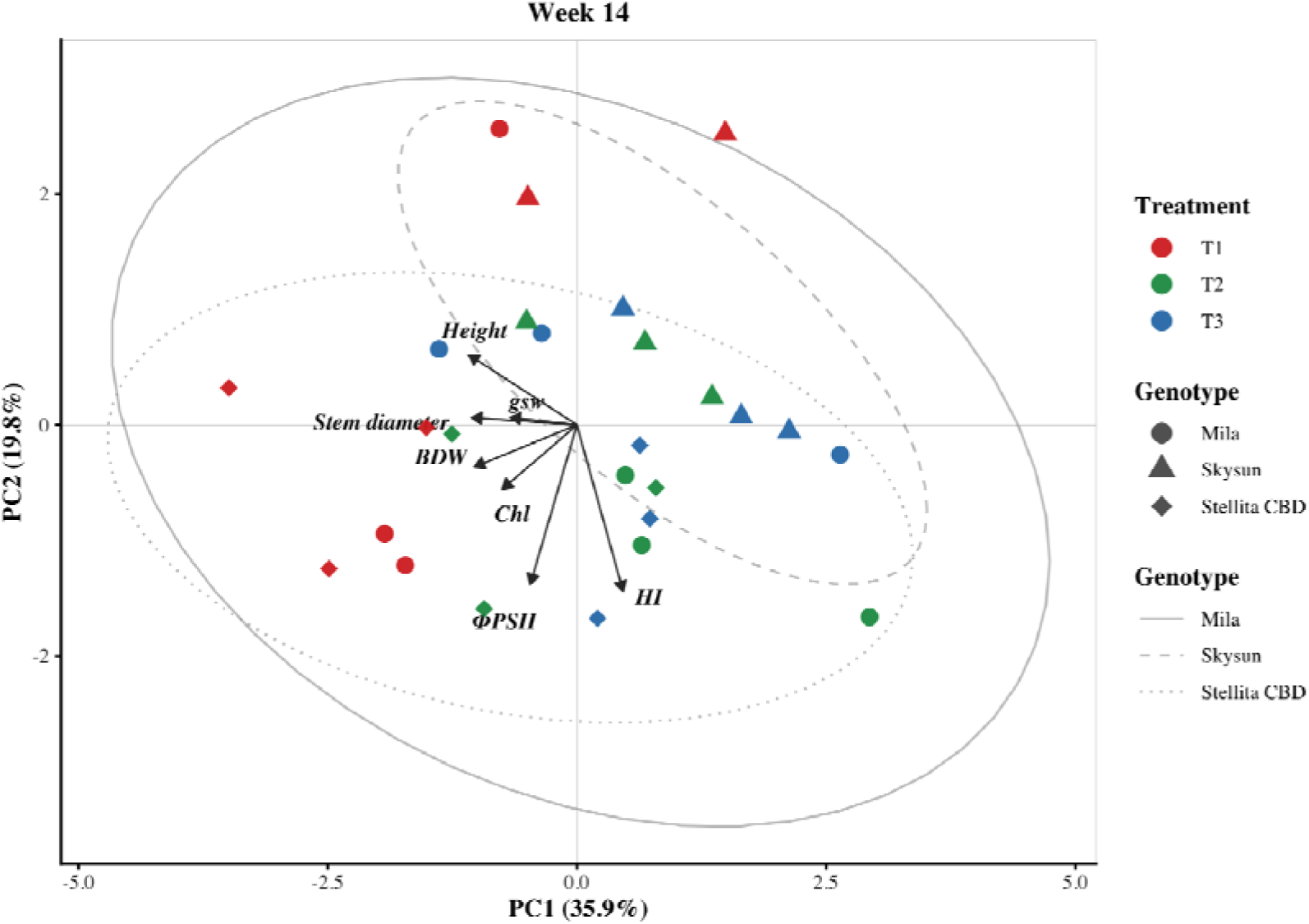
Principal component analysis (PCA) of morphological, physiological, and productivity traits of Cannabis sativa L. at week 14. Each point represents an individual plant. Point color indicates the photoperiod treatment (T1: 18 h photoperiod control; T2: 12 h light + 1 h continuous night interruption; T3: 12 h light + four 15-min pulsed night interruptions), whereas point shape identifies the genotype. Gray ellipses represent the 95% confidence regions for each genotype, with line types distinguishing genotypes (solid: MI; dashed: SS; dotted: ST). Vectors indicate the direction and relative contribution of each variable to PC1 and PC2 (Height = plant height; Stem diameter = stem diameter; Chl = total chlorophyll; ΦPSII = effective quantum yield of photosystem II; gsw = stomatal conductance; BDW = bud dry weight; HI = harvest index).

Permutational multivariate analysis of variance (PERMANOVA) confirmed significant multivariate structuring throughout the experiment. Photoperiod treatment explained the largest proportion of variation at week 8 (R² = 0.268, P < 0.001), exceeding the effect of genotype at this stage. At weeks 11 and 14, genotype and treatment contributed similarly to the observed multivariate variation (genotype R² ≈ 0.20; treatment R² ≈ 0.18). A significant genotype × treatment interaction was detected only at week 8 (P = 0.014), indicating that treatment-dependent multivariate divergence was greatest during the vegetative-to-reproductive transition.

## 4. Discussion

This study demonstrates that the effect of night-break lighting in cannabis growth and development depends not only on the total duration of nocturnal illumination but also on its temporal distribution, and that this response is strongly modulated by the genotype. Under short-day conditions, where supplemental lighting is required to maintain vegetative growth and detain inflorescence formation, these findings indicate that photoperiod manipulation can be optimized through the temporal architecture of the light signal rather than by increasing light input alone.

### 4.1 Temporal architecture of night-break signals determines photoperiodic responses

The two night-break (NB) regimes evaluated in this study were not functionally equivalent. Whereas the continuous 1-h NB treatment (T2) produced responses comparable to the commercial day-extension photoperiod (T1), the pulsed NB treatment (T3) consistently accelerated phenological progression, reduced vegetative growth and altered plant architecture in a genotype-dependent manner. Importantly, these developmental changes did not result in consistent reductions in floral biomass but rather reflected a reorganization of the plant functional phenotype. These findings agree with previous studies (Whipker et al. 2020; Gonzalez-Vanegas et al. 2024), while providing an alternative capable of reducing supplemental lighting energy consumption by approximately 27.8% (≈150 kWh) under commercial production conditions (Grover et al. 2025). Multivariate analysis clusters together T1 and T2, whereas T3 generates a different architecture in a different time frame, indicating that these lighting regimes represent functionally distinct photoperiodic signals.

Unlike the reductions in floral biomass reported under indoor conditions by Grover *et al*. (2025), no consistent yield penalties were observed in the present study. This difference is likely explained by contrasting light environments. Whereas Grover and colleagues reduced the photosynthetic daily light integral (DLI) by approximately 30%, the present experiment was conducted under greenhouse conditions where natural daylight dominated total DLI and the low-intensity NB lighting (<10 μmol m ² s ¹) functioned primarily as a photoperiodic rather than an energetic signal. Low-intensity nocturnal illumination modulates morphology and plant development (Abdulaziz, 2019; Xiao et al., 2025).

The contrasting responses observed between T2 and T3 may be explained by differences in how each lighting regime is integrated by the circadian system. The circadian clock gates responses to light–dark transitions during discrete windows of photoperiodic sensitivity (Apelt et al. 2017). Within this framework, the continuous light signal provided by T2 would be expected to coincide more consistently with these sensitive phases, thereby functioning similarly to a conventional day-extension treatment. In contrast, the discontinuous pulses delivered under T3 may only partially overlap with these windows, generating a “weak long day” due to a weaker photoperiodic signal, sufficient to prevent the development of floral clusters, yet insufficient to sustain the vegetative growth characteristic of T1 and T2.

A partial coincidence between the light pulses and circadian sensitivity windows could therefore result in an incomplete maintenance of the vegetative developmental program. This interpretation is consistent with the biology of cannabis as an annual short-day species, in which vegetative growth and reproduction compete for a finite pool of resources (Bazzaz et al. 1987; Dang et al. 2022), and where the perceived duration of the growing season influences reproductive allocation (Cohen 1976; Lundgren and Des Marais 2020).

### 4.2 Genotype-dependent sensitivity to night-break signals

The same NB regime elicited contrasting responses among the evaluated genotypes, indicating that sensitivity to photoperiodic signals is not uniform within cannabis. Genotype acted as a functional filter modulating how nocturnal light cues were integrated, a pattern supported by the multivariate analyses in which genotype explained a proportion of the observed variation comparable to that of photoperiod treatment. Consequently, a given NB regime did not produce uniform developmental responses across genotypes. Instead, the observed gradient, with Skysun and Stellita CBD exhibiting stronger responses than Mila, suggests that sensitivity to nocturnal photoperiodic signals may be more closely associated with plant architecture, developmental strategy and functional traits than with chemotype alone.

The accelerated phenological progression observed under T3 further supports this interpretation. Although all genotypes responded to pulsed NB, the magnitude of the response differed markedly, with Skysun and Stellita CBD reaching reproductive stages 17–21 days earlier than the control, whereas Mila exhibited a more attenuated response. These findings agree with previous reports describing substantial genetic variation in photoperiod sensitivity in cannabis, where taller genotypes generally exhibit stronger photoperiodic responses than more compact plants (Zhang et al. 2021). The reduced sensitivity of Mila, the most compact genotype evaluated, suggests that responsiveness to NB may reflect broader functional differences among genotypes, potentially related to breeding history or cultivation group.

The earlier phenological progression and architectural reorganization observed under T3 are consistent with adaptive responses described for annual species experiencing a shorter or less predictable effective growing season. Under such conditions, plants tend to prioritize reproductive development over prolonged vegetative growth, thereby maximizing the probability of completing their life cycle without necessarily experiencing physiological stress (Franks et al. 2007; Hansen et al. 2013). Within this framework, the genotype-dependent responses observed here suggest that NB treatments may modify developmental scheduling according to each genotype’s intrinsic sensitivity to photoperiodic cues. Together, these findings highlight genotype as a major determinant of NB effectiveness and emphasize the importance of considering genetic background when designing photoperiod management strategies for cannabis cultivation.

### 4.3 Developmental reprogramming without physiological impairment

The genotype-dependent sensitivity to NB treatments was also reflected in plant growth. The reduced height of T3 was accompanied by a faster shortening of the apical internodes, particularly in Skysun, where weekly shortening rates reached 31–35% compared with 15% in T1. Progressive internode shortening is recognized as one of the earliest indicators of reproductive transition in cannabis (Spitzer-Rimon et al. 2019). Rather than reflecting growth inhibition, these responses indicate an architectural reorganization of the plant. More compact canopies do not necessarily compromise productivity and may even improve light interception and biomass distribution within the canopy while reducing the need for intensive pruning practices (Danziger and Bernstein 2021; Carlson et al. 2021).

NB treatments also modified the temporal dynamics of growth. Beyond plant height, the most sensitive phenotypes modified growth stabilization timing, which indicates that the temporal distribution of nocturnal light reprogrammed developmental trajectories rather than simply reducing vegetative growth (Vanhove et al. 2017; Stack et al. 2021; Dang et al. 2022).

The transient increase in total chlorophyll content observed under T3 around week 11 for all genotypes coincided with the transition toward advanced reproductive stages, during which pigment concentrations may increase before the onset of senescence (Lu et al. 2002; Falqueto et al. 2010). In the present study, the subsequent decline in chlorophyll content was not accompanied by a reduction in Φ_PSII_, consistent with phased models of leaf senescence in which pigment degradation precedes the loss of photosynthetic function (Wingler et al. 2004; Guo and Gan 2005). These results suggest that the chlorophyll peak observed under T3 is a hallmark of cannabis reproductive development and reflects accelerated developmental progression rather than physiological impairment. Future studies incorporating complementary markers of developmental status and senescence, including metabolic profiling, could help to validate this interpretation.

### 4.4 Productivity and quality: biomass partitioning and resource allocation

The architectural reorganization described above becomes particularly relevant when considered in the context of biomass allocation and productivity. In cannabis, plant architecture—including traits such as height, stem diameter and internode length—is closely associated with the partitioning of assimilates between vegetative growth and reproduction, ultimately determining the efficiency with which biomass is converted into harvestable yield (Naim-Feil et al. 2021; Danziger and Bernstein 2021; Carlson et al. 2021). Within this framework, reduced plant stature or canopy compaction does not necessarily imply lower productivity but may instead reflect a shift in biomass allocation toward reproductive tissues. This interpretation is consistent with annual photoperiod-sensitive species, in which accelerated developmental schedules are commonly accompanied by reduced investment in vegetative structures and increased reproductive allocation (Bazzaz et al. 1987; Dang et al. 2022).

Despite the phenological and architectural modifications induced by NB treatments, floral dry biomass (BDW) remained largely unchanged across treatments in most genotypes, indicating that developmental reprogramming did not compromise reproductive yield. Instead, BDW was primarily determined by genotype, confirming that genetic background remained the main driver of productivity (Sullivan et al. 2025). In contrast, harvest index (HI) was consistently more responsive to photoperiod manipulation, increasing by 15–64% under NB treatments. These results indicate that NB primarily altered biomass partitioning rather than total biomass production, promoting greater biomass allocation to reproductive tissues. Similar plasticity in HI has previously been described in cannabis (Naim-Feil et al. 2021).

This functional trade-off is further supported by the multivariate analysis at harvest, where HI and structural traits (plant height and stem diameter) were associated with opposite directions along the principal component space. Such patterns are consistent with the classical trade-off between structural investment and reproductive allocation described for annual species (Lundgren and Des Marais 2020; Burgel et al. 2020; Carlson et al. 2021).

Cannabinoid composition remained primarily genotype-dependent, with NB acting as a secondary environmental modulator. Mila maintained stable CBD concentrations across treatments, whereas Skysun and Stellita CBD exhibited genotype-specific responses. Such patterns agree with previous studies demonstrating that cannabinoid composition is largely determined by genetic regulation, while environmental conditions exert a comparatively smaller influence (Burgel et al. 2020; Rashid et al. 2021). Likewise, photoperiod manipulation during flowering has been shown to increase CBD yield in CBD-dominant chemotypes, whereas the same photoperiodic changes can increase floral biomass while reducing THC concentration in THC-rich genotypes, highlighting a chemotype-dependent response (Peterswald et al. 2023). Consistent with these observations, the present study indicates that low-intensity NB, acting as a photoperiodic and circadian regulator rather than as an additional source of photosynthetic radiation, induces genotype-specific adjustments in cannabinoid accumulation instead of uniform changes across genotypes.

## 5. Conclusions

The night-break (NB) regimes evaluated in this study effectively delayed reproductive development in cannabis to a degree comparable with the conventional day-extension photoperiod. However, the two NB strategies were not functionally equivalent, demonstrating that the biological effectiveness of nocturnal light depends not only on its total duration but also on its temporal organization. Plant responses were strongly genotype-dependent, with genotype acting as a functional filter that modulated the integration of photoperiodic signals. The pulsed NB treatment (T3) accelerated developmental progression and promoted a more compact plant architecture by reprogramming developmental trajectories rather than altering intrinsic growth rates. These developmental changes occurred without evidence of physiological impairment and were accompanied by changes in biomass partitioning that increased harvest index while largely maintaining floral biomass and cannabinoid composition. Collectively, these findings demonstrate that NB photoperiods represent a viable and energy-efficient light management strategy whose implementation should be optimized according to genotype, production objectives, and the temporal architecture of the nocturnal light signal.

## Acknowledgements

The authors acknowledge the Agencia Distrital para la Educación Superior, la Ciencia y la Tecnología (ATENEA) for funding the project “Interrupción de la noche como alternativa a la extensión de la fase diurna para producción de cannabinoides” through Contract No. 289-2024, under which this research was conducted. We thank PANACEA CL S.A.S. for providing greenhouse facilities, technical support and logistical assistance during the experimental phase carried out in La Vega, Cundinamarca, Colombia. We also acknowledge the Plant Physiology Laboratory at the Universidad Nacional de Colombia for their assistance with data processing.

## CRediT authorship contribution statement

Rodríguez-Roncancio, Maria Paula: Conceptualization, Methodology, Investigation, Data curation, Formal analysis, Visualization, Writing – original draft, Writing – review & editing. Castillo-Hernández, Laura Camila: Investigation, Data curation. Love González-Sarmiento, Nitro: Investigation. Luz Marina Melgarejo: Conceptualization, Methodology, Supervision, Resources, Writing – review & editing. Felipe Sarmiento: Conceptualization, Methodology, Supervision, Project administration, Funding acquisition, Resources, Writing – review & editing.

## Data availability

The data supporting the findings of this study are available from the corresponding author upon reasonable request.

## Ethics approval

Ethics approval was not required in this study as only plant material was used in the experiments.

