## supplementary material for "Night-break photoperiod timing shapes growth trajectories, functional phenotype, and biomass partitioning in *Cannabis sativa* L. in a genotype-dependent manner"

**Acta Physiologiae Plantarum**

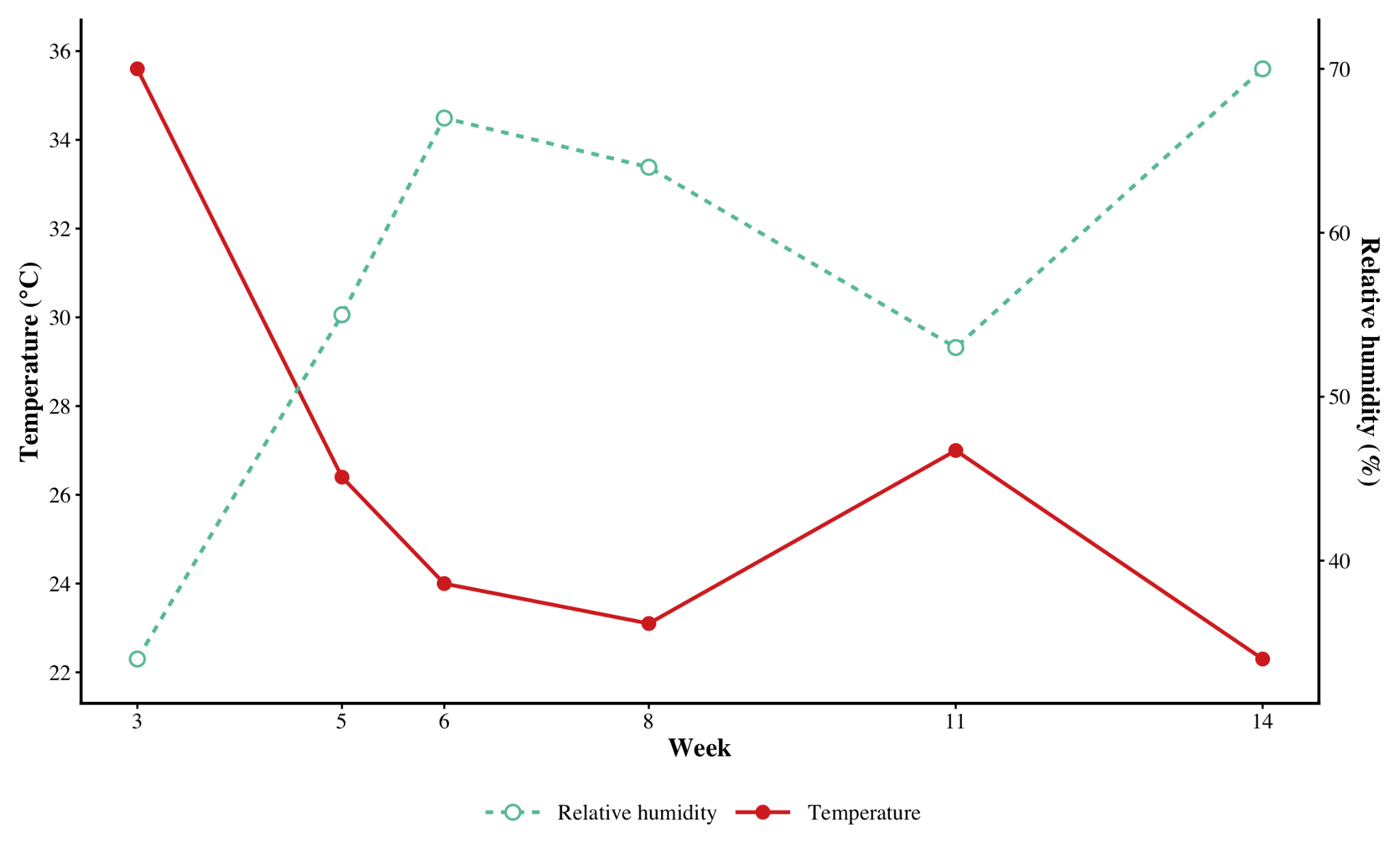

**Supplementary Figure S1.** Air temperature (°C) and relative humidity (%) recorded at each sampling date (weeks 3–14) during the experimental period. Relative humidity is displayed on the secondary (right) axis to facilitate comparison with temperature.

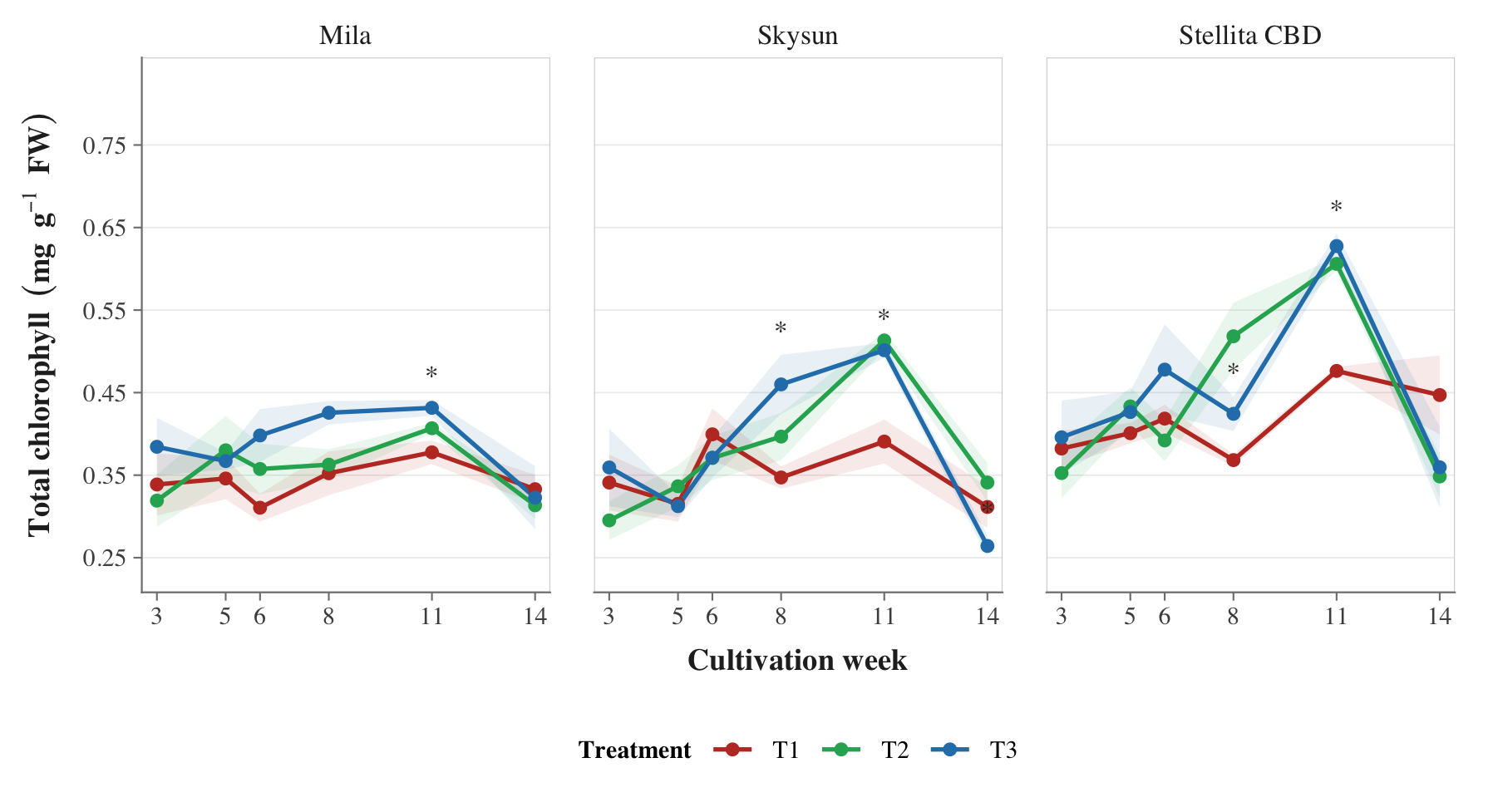

**Supplementary Figure S2.** Temporal changes in total chlorophyll content of three *Cannabis sativa* genotypes under three photoperiod treatments: T1 (commercial day-extension control), T2 (1 h continuous night-break), and T3 (4 × 15 min pulsed night-break). Symbols represent treatment means and shaded areas indicate the standard error (SE). Asterisks denote sampling weeks in which significant differences among treatments were detected within each genotype (*P* < 0.05).

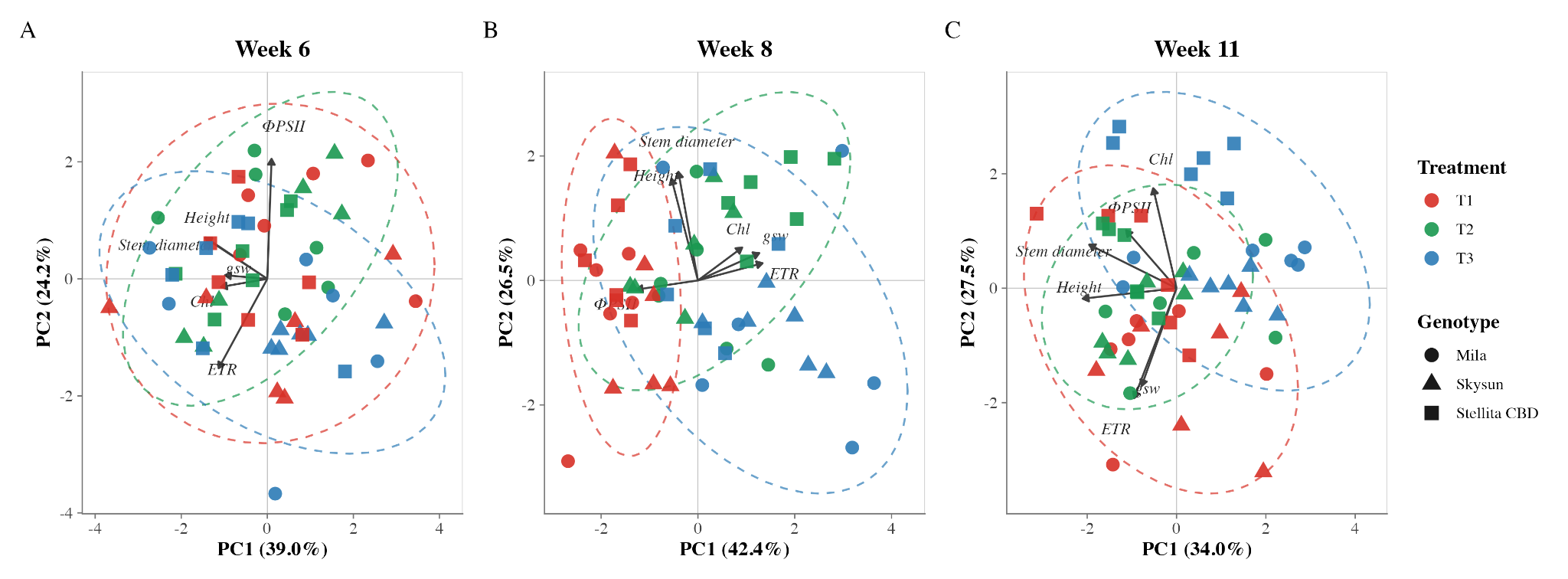

**Supplementary Figure S3.** Principal component analyses (PCA) of morphological and physiological traits at weeks 6 (A), 8 (B), and 11 (C) in three *Cannabis sativa* genotypes grown under three photoperiod treatments. Each point represents an individual plant. Point color indicates the photoperiod treatment (T1, T2 and T3), whereas point shape identifies the genotype (Mila, Skysun and Stellita CBD). Dashed ellipses represent the 95% confidence regions for each treatment. Vectors indicate the direction and relative contribution of each variable to the first two principal components

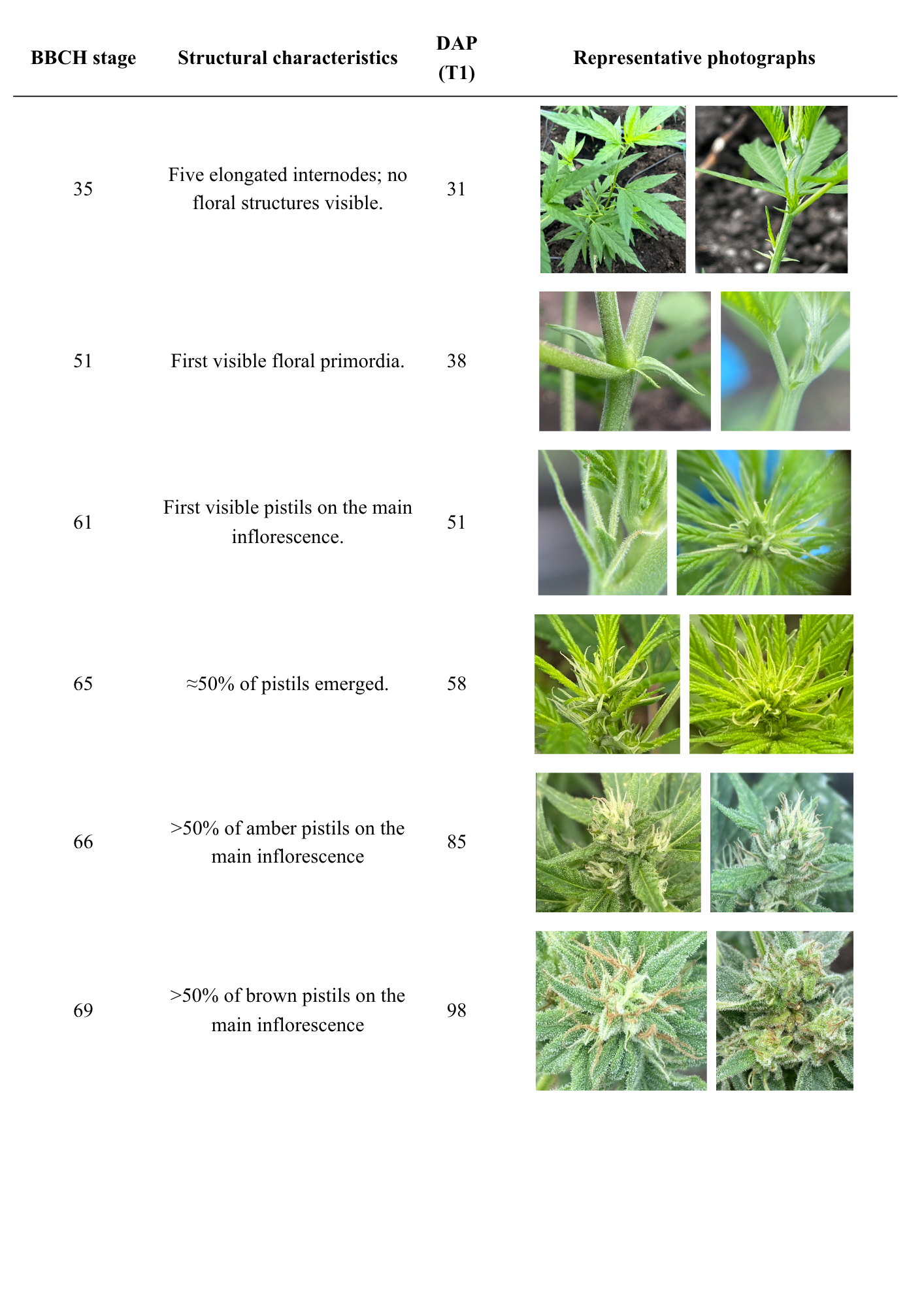

**Supplementary Figure S4.** Representative BBCH phenological stages used to characterize reproductive development in Cannabis sativa. The figure summarizes the diagnostic structural characteristics and representative photographs used to assign each developmental stage. Approximate days after planting (DAP) correspond to the commercial day-extension treatment (T1) and are provided as a reference for stage identification.

**Supplementary Table S1.** Nonlinear models used to describe plant growth trajectories in *Cannabis sativa* under different photoperiod treatments. The table summarizes the equations, model parameters, and their biological interpretation for plant height, stem diameter, and apical internode length.

| **Variable** | **Model** | **Parameters** | **Biological meaning** |
| --- | --- | --- | --- |
| Plant height | $Height(t) = \frac{Asym}{1+ exp(\frac{xmid - t}{scal}) + \varepsilon}$ | H(t): predicted plant height at cultivation week t; Asym: estimated asymptotic maximum; xmid: week at which plants reach 50% of the estimated maximum height; scal: scale parameter related to the slope around the inflection point. | Describes sigmoidal plant height growth over time. Asym represents the estimated maximum height, xmid identifies the time at which 50% of the asymptotic height is reached, and scal determines the steepness of the growth trajectory around the inflection point. |
| Stem diameter | $Diameter (t) = \frac{Asym \cdot t}{K - t} + \varepsilon$ | D(t): stem diameter at cultivation week t; Asym: estimated asymptotic maximum stem diameter; K: half-saturation constant, corresponding to the week at which plants reach 50% of the estimated maximum diameter. | Describes asymptotic stem diameter expansion. Asym represents the estimated maximum stem diameter, whereas K indicates how rapidly plants approach the asymptotic diameter. |
| Apical internode length | $Internode (t) = a\cdot e^{-bt}+\varepsilon$ | I(t): apical internode length at cultivation week t; a: estimated initial internode length; b: exponential rate of internode shortening over time. | Describes the progressive shortening of the apical internode. The parameter a represents the initial estimated internode length, whereas b indicates the rate at which internode length decreases during plant development. |

**Supplementary Table S2.** Days after planting (DAP) required to reach key phenological stages (BBCH 51, 61, and 69) in three Cannabis sativa genotypes grown under three photoperiod treatments. Values are presented as the median with the interquartile range (IQR; Q1–Q3). Numbers in parentheses indicate the proportion of plants reaching each stage (n/N). BBCH 51, first visible floral primordia; BBCH 61, first visible pistils on the main inflorescence; BBCH 69, >50% of pistils senescent. DAP values were constrained by the sampling schedule (21, 35, 42, 56, 77, and 98 days after planting).

| **BBCH stage** | **Mila** | | | **Skysun** | | | **Stellita CBD** | | |
| --- | --- | --- | --- | --- | --- | --- | --- | --- | --- |
|  | T1 | T2 | T3 | T1 | T2 | T3 | T1 | T2 | T3 |
| **51** | 28 [21–35] (6/6) | 21 [21–32] (6/6) | 21 [21–21] (6/6) | 46 [35–56] (6/6) | 35 [35–35] (6/6) | 28 [21–35] (6/6) | 35 [35–35] (6/6) | 35 [35–35] (6/6) | 21 [21–32] (6/6) |
| **61** | 35 [35–35] (6/6) | 21 [21–32] (6/6) | 21 [21–21] (6/6) | 56 [56–56] (6/6) | 56 [56–56] (6/6) | 38 [35–52] (6/6) | 56 [56–56] (6/6) | 56 [56–56] (6/6) | 56 [40–56] (6/6) |
| **69** | 98 [98–98] (3/6) | 98 [82–98] (6/6) | 77 [77–93] (6/6) | 98 [98–98] (3/6) | 98 [93–98] (4/6) | 77 [77–93] (6/6) | 98 [98–98] (1/6) | 98 [98–98] (3/6) | 98 [98–98] (6/6) |
